# Artificial Intelligence Models for Classifying Wrist Ligament Injuries Using Synthetically-Generated Joint Proximity Maps from Finite Element Models

**DOI:** 10.64898/2026.06.17.733030

**Authors:** Hsuan-Yu Chen, Jon Camp, Taylor P. Trentadue, Andrew R. Thoreson, Shuai Leng, David R. Holmes, Sanjeev Kakar, Kai-Nan An, Kristin D. Zhao, Thor E. Andreassen

## Abstract

**Background/Purpose:** Diagnosing wrist ligament injuries is challenging; early detection and treatment are important to prevent osteoarthritis progression. Interosseous proximity maps, a proxy measure for joint space, can be generated from volumetric imaging data and may provide important information about wrist health. Artificial intelligence (AI) could enhance accuracy of noninvasive diagnosis based on imaging-derived metrics. This work demonstrates feasibility of AI training using synthetic proximity map data generated from finite element models (FEMs).

**Methods:** Personalized wrist FEMs for two asymptomatic participants were created from four-dimensional computed tomography-derived anatomic and kinematic data. Monte Carlo sampling varied 22 ligament material properties and simulated 7,500 unique injury scenarios generating 9,000,000 labeled red, green, and blue (RGB) images of interosseous proximity vector fields from FEM-derived motions. Images were associated with 17 descriptive metrics, including gross wrist angles and bone surface pairs, and used to develop mixed-input convolutional neural networks (CNNs). Model performance was evaluated for identifying specific ligament injuries.

**Results:** Average area under receiver operating characteristic curve (AUROC) for CNNs was 0.757 across all injury types and kinematics. In a subset with clinically-relevant functional angles, the average AUROC was 0.824. Best-performing individual ligament AUROCs ranged from 0.807 to 0.999. Sensitivities and specificities exceeded 0.99 for some ligament injury simulations under specific wrist angles and bone surface pairs.

**Conclusion:** This study demonstrates the feasibility of using synthetic data from FEMs to train AI models for classifying wrist ligament injuries. Proximity-based RGB images may be a relevant biomarker of ligamentous injury.

## Introduction

Wrist ligament injuries are common, with prevalence varying by the mechanism of injury and demographic under study (1). Common wrist injuries, such as scapholunate interosseous ligament (SLIL) tears (Figure 1), may create carpal instability and have been reported in approximately 20-32% of distal radius fractures, with the highest incidence observed in intra-articular fracture patterns (2). Up to 44% of patients sustaining fall-on-outstretched-hand (FOOSH) injuries may develop carpal instability within 24 months (3). These injuries may predispose patients to developing osteoarthritis through disrupted carpal kinematics, articular malalignment, and altered load distribution (4, 5). Progression from ligament injury to static malalignment and osteoarthritis contribute to substantial morbidity, including chronic pain, functional impairment, and increased healthcare utilization (6).

**Figure 1:**
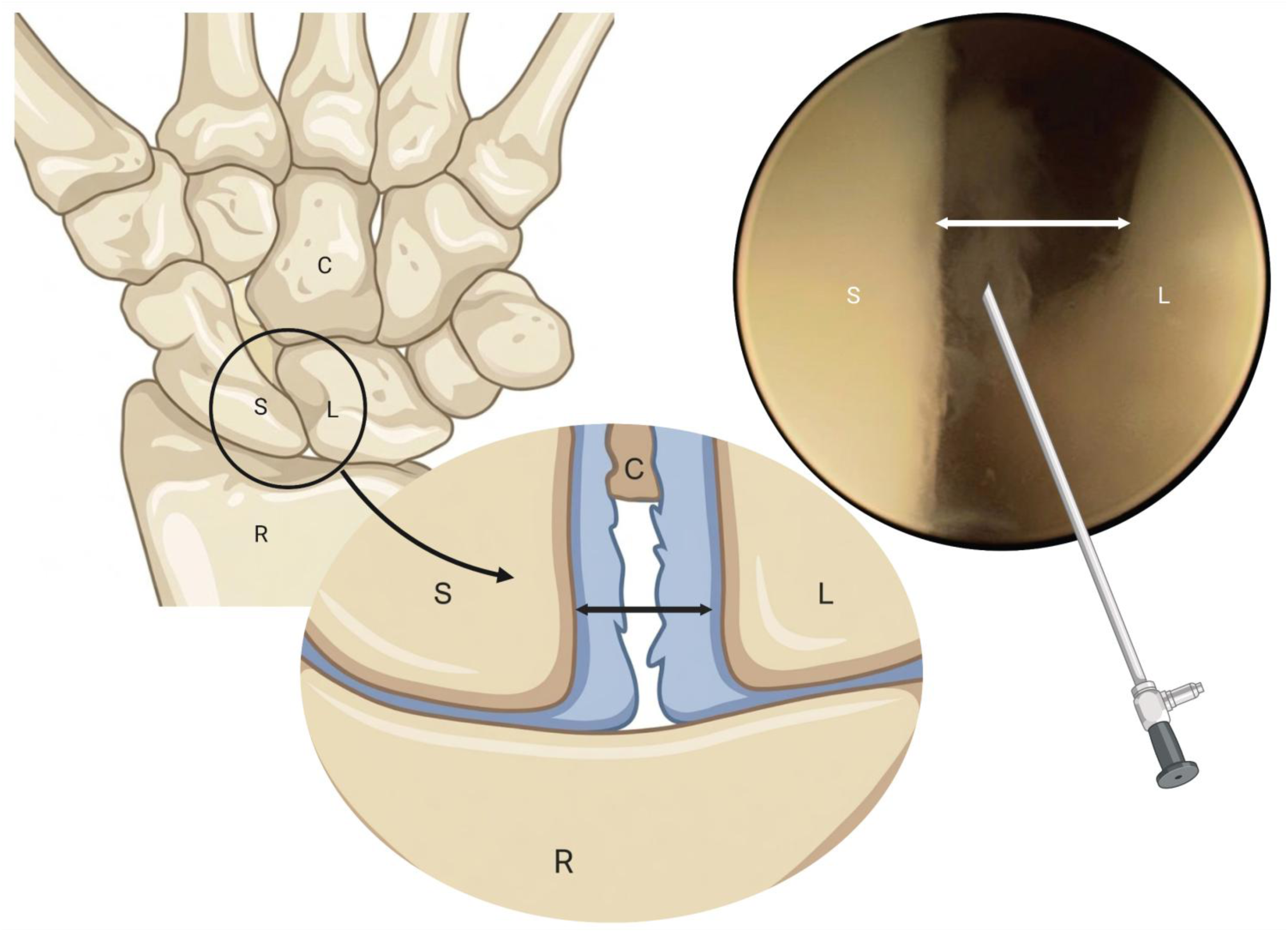
Schematic depiction of scapholunate ligament injury and representative arthroscopic visualization demonstrating the separation between carpal bones under stress. C: capitate, L: lunate, R: radius, S: scaphoid. (Created in BioRender. Chen, HY. (2026) https://BioRender.com/vwrxkbz)

Diagnosing wrist ligament injuries involve physical examination, routine and advanced imaging, and arthroscopic evaluation (Figure 1). Equivocal or unremarkable examination findings may delay treatment of wrist ligament injuries (6, 7). Routine clinical assessments, including patient histories and physical examination findings alone, yield low negative predictive values (8). Conventional radiographs and ultrasound often serve as initial imaging modalities for querying carpal alignment or ligament injury. Advanced imaging modalities including magnetic resonance imaging (MRI), MR arthrography, computed tomography (CT), or dynamic four-dimensional CT (4DCT) are frequently used by clinicians to stage ligamentous injuries and assess carpal instability (6, 9).

3.0T MRI yields sensitivity of 70-87% and specificity of 90-97% for SLIL tears (10). Wrist arthroscopy remains the preferred diagnostic modality due to its high conclusive reliability, as negative MRI findings cannot exclude clinically significant injuries (11–13). Arthroscopy may facilitate direct visualization of ligament injury, offering critical information for standardized classification systems that guide surgical decision-making (14, 15). Still, it remains an invasive procedure requiring specialized training, equipment, and surgical expertise. Moreover, wrist stability relies on the intricate interplay of multiple ligaments; disruption of these structures initiates a cascade of biomechanical alterations leading to progressive derangements in loading patterns and cartilage degeneration (5). While imaging can often identify the presence of injury, it remains difficult to identify the specific permutations of ligaments injured. This nuance is clinically relevant and an area of active research (16).

Finite element models (FEMs) are frequently applied in engineering settings, with growing applications in biomechanics and orthopaedics (17). Orthopaedics-oriented FEMs are typically achieved by deriving geometric structures from volumetric imaging, such as CT or MRI (18). FEMs can offer insights into the complex interactions between bones, ligaments, cartilage, muscles, and tendons that would be difficult with other methods given the inherent complexity and number of structures involved. Once developed, these models provide valuable quantitative metrics that are often impractical to acquire *in vivo* including contact pressures, stress distributions, ligament forces, and joint kinematics(18, 19).

Our prior work utilizing 4DCT has captured individuals with and without ligament injuries to investigate changes to arthrokinematics via image-derived joint proximity maps (interosseous bone distances) (20, 21). Moreover, we have previously developed individualized wrist models and demonstrated the feasibility of predicting joint contact mechanics (22) and a clinically-relevant provocative maneuver, namely a scaphoid shift maneuver to isolate scapholunate ligament tears (23). Still, the complexities of patient-specific joint morphology, material properties, ligamentous and tendinous attachment sites, and boundary conditions limit clinical translation of FEMs (24). Additionally, the time required to generate patient-specific FEMs is a key barrier (25).

Artificial intelligence (AI) can rapidly classify data to support routine clinical decision-making. Models trained using biomechanical data from musculoskeletal simulations have demonstrated success in classifying wrist pathologies (26). However, AI models with limited complexity struggle to identify the presence of injuries, particularly when accounting for individual variability (20). While more sophisticated AI models will likely improve diagnostic performance, this requires massive amounts of existing data to train these models, which currently remains scarce (27).

Synthetically-generated data derived from FEMs could provide comprehensive training datasets for AI models by offering larger datasets than what can be collected via physical testing and clinical data alone. Models trained on such simulated data may also better identify subtle features associated with early-stage disease. Furthermore, the synergy offered by FEMs and AI may enhance understanding of injury mechanisms by identifying biomechanical features that differentiate pathological states.

The objective of this study was to develop and evaluate an AI model to classify clinically-relevant, simulated wrist ligament injuries trained with joint proximity maps derived from two personalized wrist FEMs. Synthetic proximity map images of interest were generated under controlled ligament injury conditions and used to train and test the AI classifiers. We hypothesize that specific combination of wrist joint kinematic features and articular bone surface pairings will yield improved AI diagnostic performance for detecting various types of wrist ligament injuries over previously reported methods.

## Materials and Methods

The overall strategy for this work is provided in Figure 2, demonstrating the three main phases: FEM development, synthetic dataset generation, and AI model development.

**Figure 2:**
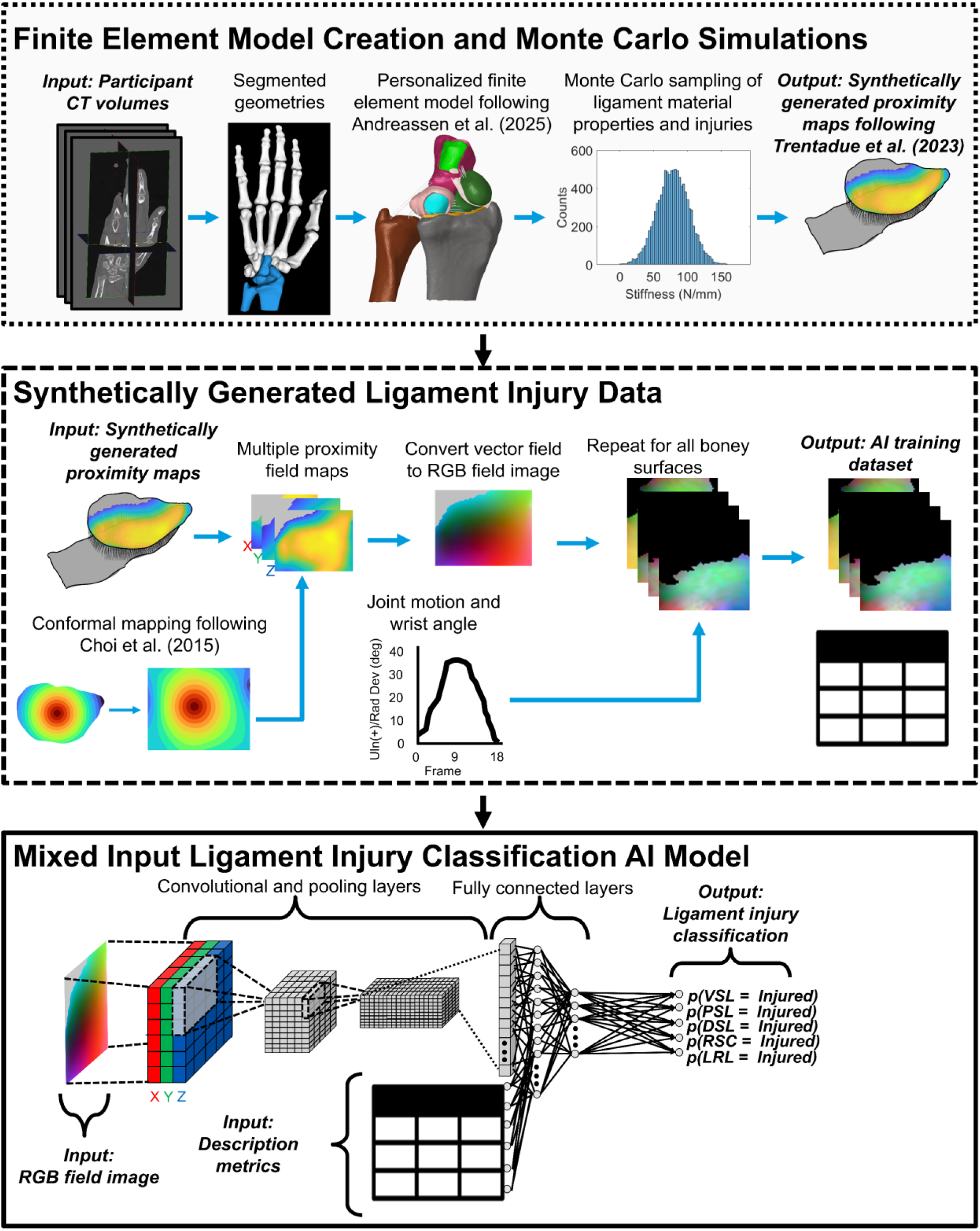
Overall Workflow. (Dotted Border Box) Process to create personalized finite element models from experimental static CT and four-dimensional computed tomography (4DCT) data. Models simulate different motions under different random sets of ligament material properties and injury conditions. Predicted motions are used to calculate interosseous joint proximity maps, a representation of joint arthrokinematics. (Dashed Border Box) Steps to convert simulated proximity maps into machine-readable consistent red-green-blue (RGB) vector field images and descriptive metrics of simulation including radiocapitate angle, scale of proximity vectors, and specific bone surface pair. (Solid Border Box) AI classification model using the mixed inputs (images and descriptive metrics) to predict presence of ligament injury in each ligament separately. This work combined code from several previous studies including prior modeling work from Andreassen et al. (2025), prior proximity map generation code from Trentadue et al., (2023), and prior conformal mapping code from Choi et al., (2015) (21, 22, 33).

Following IRB approval (NCT 04736537) and written informed consent, static and 4DCT data were collected from a normative cohort of asymptomatic participants (28, 29). Static CT data were collected with participants in an anatomically-neutral position. Dynamic data were collected using a previously-described 4DCT protocol during wrist flexion-extension or radioulnar deviation (20). Carpal bones were segmented from static CT images and registered to corresponding positions in each dynamic image volume (30) to obtain carpal geometries and kinematics, respectively. These data were introduced to a previously-described workflow to develop two personalized wrist FEMs (22). Cartilage and ligament attachment sites were predicted using a combination of algorithmic and non-linear morphing techniques (Figure 2) (31).

The wrist FEMs are versatile, allowing for variations of ligament material properties, namely stiffness and reference strain (ligament initially slack or taut), and simulation of injuries to the ligament fibers. FEMs were moved through wrist flexion-extension or radioulnar deviation arcs by applying 4DCT-derived rotations to the capitate (32), with motion of other bones driven by residual forces in the modeled ligaments and contact mechanics between adjacent bodies. Bone osteokinematics (rotations and translations) were captured over the range of simulated motion and then used to calculate interosseous joint proximity maps. This process was repeated for different permutations of ligament material properties, joint motions, and ligament injury states to develop training data for the AI model. Some permutations resulted in proximities that did not meet a minimum distance threshold of 5mm or an angular threshold of within 60 degrees and were thus excluded.

The prior proximity maps are information-rich and have previously been shown to detect subtle changes resulting from injuries(20). However, the inherent spatiotemporal nature of this data, namely time-varying, non-uniform 3D vector fields, poses significant challenges for integration into AI. Consequently, only a limited number of approaches can handle this data effectively. In contrast, images, both 2D and 3D, are widely used as standard inputs in AI. Many tools have been developed to utilize this data. Therefore, a significant portion of this work focused on developing methods to convert proximity maps into image formats that AI models can utilize more efficiently.

A novel procedure was used to convert the relative 3D proximity map vectors into a set of red-green-blue (RGB) vector field images (Figure 2). First, each proximity map was morphed to a template bone using the aforementioned algorithm (31) to establish a common reference surface. Next, conformal mapping techniques (33) were used to project the 3D surface vector onto a two-dimensional plane. The interosseous proximity in each anatomic direction was converted to the value of a corresponding color in the image. Ultimately, the RGB field captured radial/ulnar translation (red, x), dorsal/volar translation (green, y), and proximal/distal translation (blue, z). For each simulation, these RGB images were created for different articulating bone surface pairs and across the entire range of motion (Figure 3). The final resolution of the RGB conformal proximity map images was 64×64 pixels. In addition, to the RGB images, a set of 17 descriptive metrics was included in model training. These features include the pair of bone surfaces the image represents, the wrist motion scales, and the relative scale of the RGB images (Figure 3). The model outputs were the predicted binarized injury status (intact versus injured) for each of the five ligamentous stabilizers individually (Figure 3) resulting in 32 total permutations of ligament injuries.

**Figure 3:**
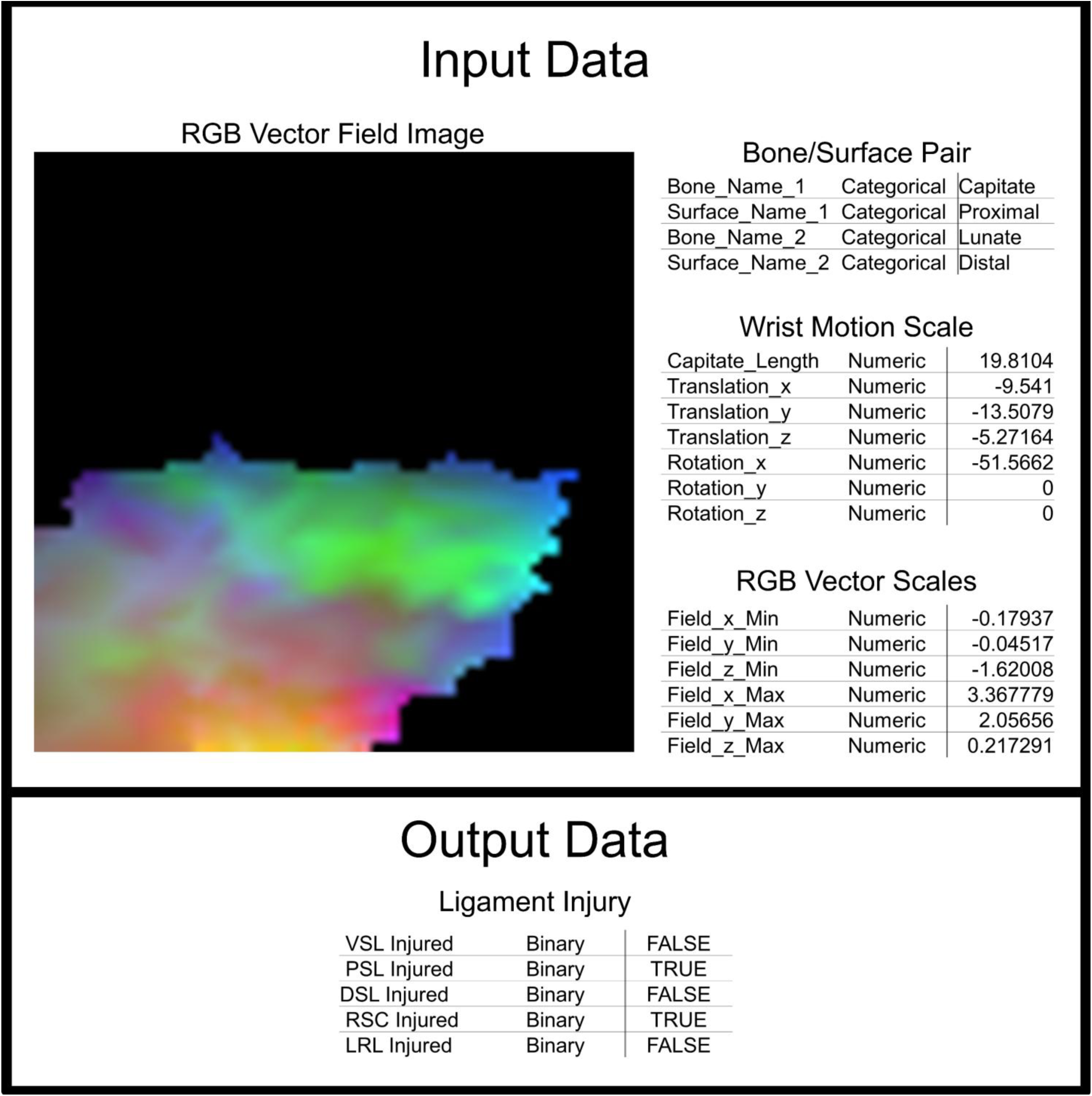
Demonstrative synthetic data used to train the AI model. Input data includes a RGB vector field image with colors representing the relative distances between the adjacent bones, as well as descriptive metrics describing the contextual information of the image. The output data are binary values for which of the five ligaments were injured for the simulation. In total the binary output data are independent of one another for the simulations, and therefore, there are 32 total permutations of ligament injuries simulated.

Using the previous modeling and conformal mapping pipeline, a Monte Carlo sampling method varied the 22 ligament material properties randomly according to physiological material property distributions derived from prior cadaveric work (22). In addition, each sample created a random permutation of injury to the five primary stabilizers of the scapholunate joint: the volar SLIL (VSL), proximal or membranous SLIL (PSL), dorsal SLIL (DSL), radioscaphocapitate ligament (RSC), and long radiolunate ligament (LRL). In each simulation, each of the five ligamentous stabilizers had a 33% chance of being injured. In other words, the probability that all ligaments were intact within a simulation was (2/3)^5^ or 16%. In total, FEMs performed over 7,500 simulations, yielding over 9,000,000 labeled RGB images and associated descriptive metrics. All images were assigned to 10 different angle bins for FE (10° increments) and 10 bins for RU (5° increments) based on the total range of motion. This was done as variation within the bins was likely insignificant, and these ranges would better approximate clinical setup in an imaging scanner where prescriptions of the wrist position within the imaging scanner would be limited to visual inspection.

Data were divided into training (80%), validation (10%), and testing (10%) portions and used to develop mixed-input, multi-label convolutional neural networks (CNNs, Figure 2). CNNs were created in TensorFlow (34). The models used an RGB image as input to a series of 2D convolutional layers before being input for a set of fully-connected dense layers (Figure 2 and Figure 4). The descriptive metrics were concatenated as input activations to the first dense layer (Figure 2 and Figure 4). Several dense layers were included before output labels were predicted. The Adam optimizer was used to minimize binary cross-entropy loss between predicted ligament injuries and the ground truth of the training dataset (35). Crucially, this allowed the model to simultaneously predict which ligaments were injured without assuming common injury progression.

**Figure 4:**
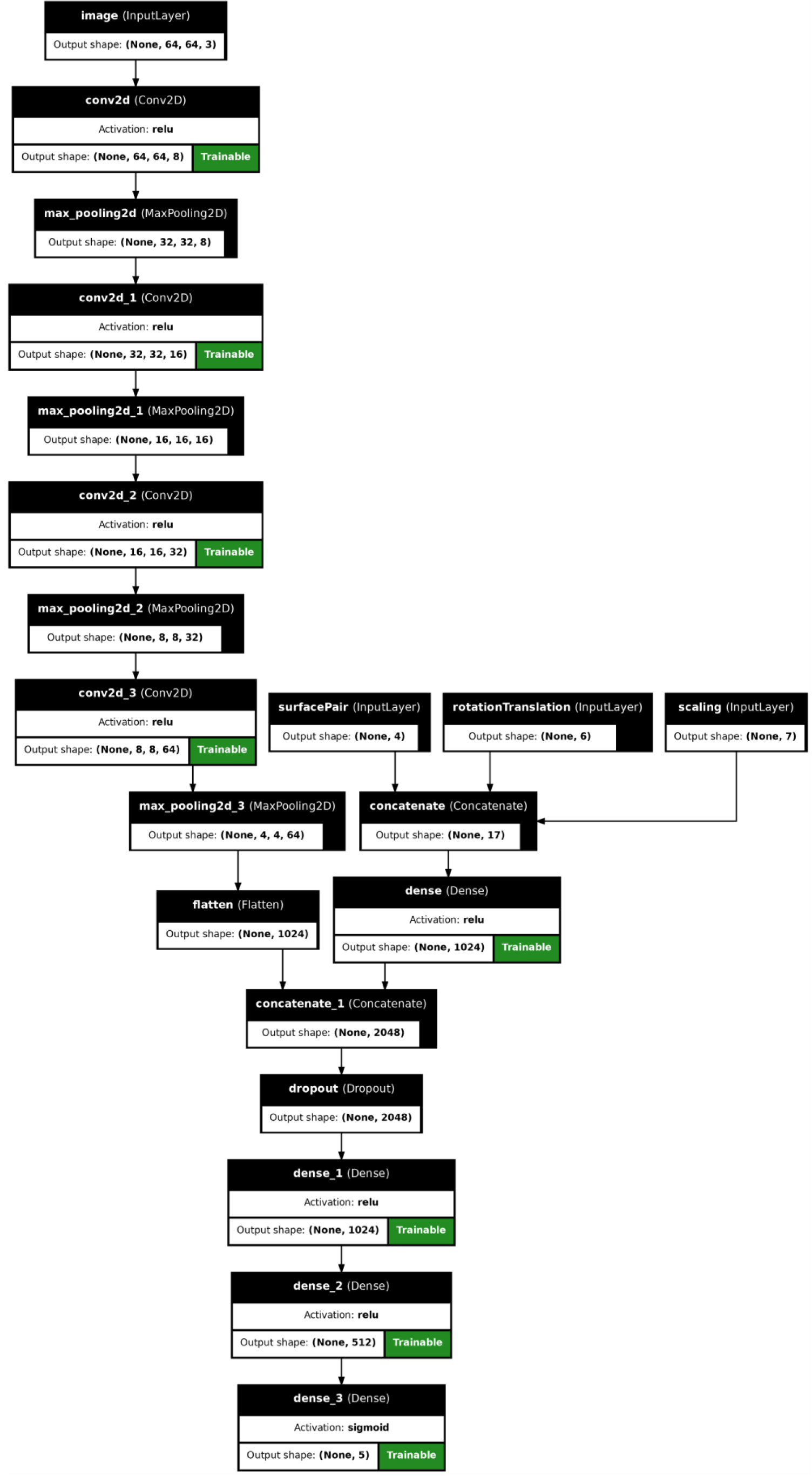
Structure of trained AI models, with mixed input data, namely bone surface conformal mapped proximity maps (RGB images) and descriptive metrics of image (Tabular data).

The validation dataset was used for tuning model hyperparameters and limit model overfitting. The testing dataset was used to report performance metrics including accuracy, sensitivity, specificity, precision, recall, F1 score, and area under the receiver operating characteristic curve (AUROC) for each ligament injury. Descriptive statistics including average, minima, and maxima were calculated for each metric across the range of subsets for each ligament based on the wrist angle range and the bone surface pair. Only subsets with at least 500 examples were included in the analysis.

## Results

When looking at identifying individual ligaments injured, CNNs achieved an overall accuracy of 0.770 and an average AUROC of 0.757 when predicting injury across all permutations, bone surface pairs, and wrist angles.

However, when grouping by different subsets of injury types, articulating bone surface pairs, and wrist angles, the range of AUROCs varied between 0.501 and 0.999 for different subsets (Table 1). As such, the remaining analyses were divided into performance metrics for different subsets. For example, in one such subset, namely datapoints with images between the proximal-side surface of the scaphoid and the radial-side surface of the capitate bones for wrist radial deviation angles between 10 - 15° (Figure 5A), the AUROC varied between 0.574 to 0.975 for the five ligaments.

**Figure 5:**
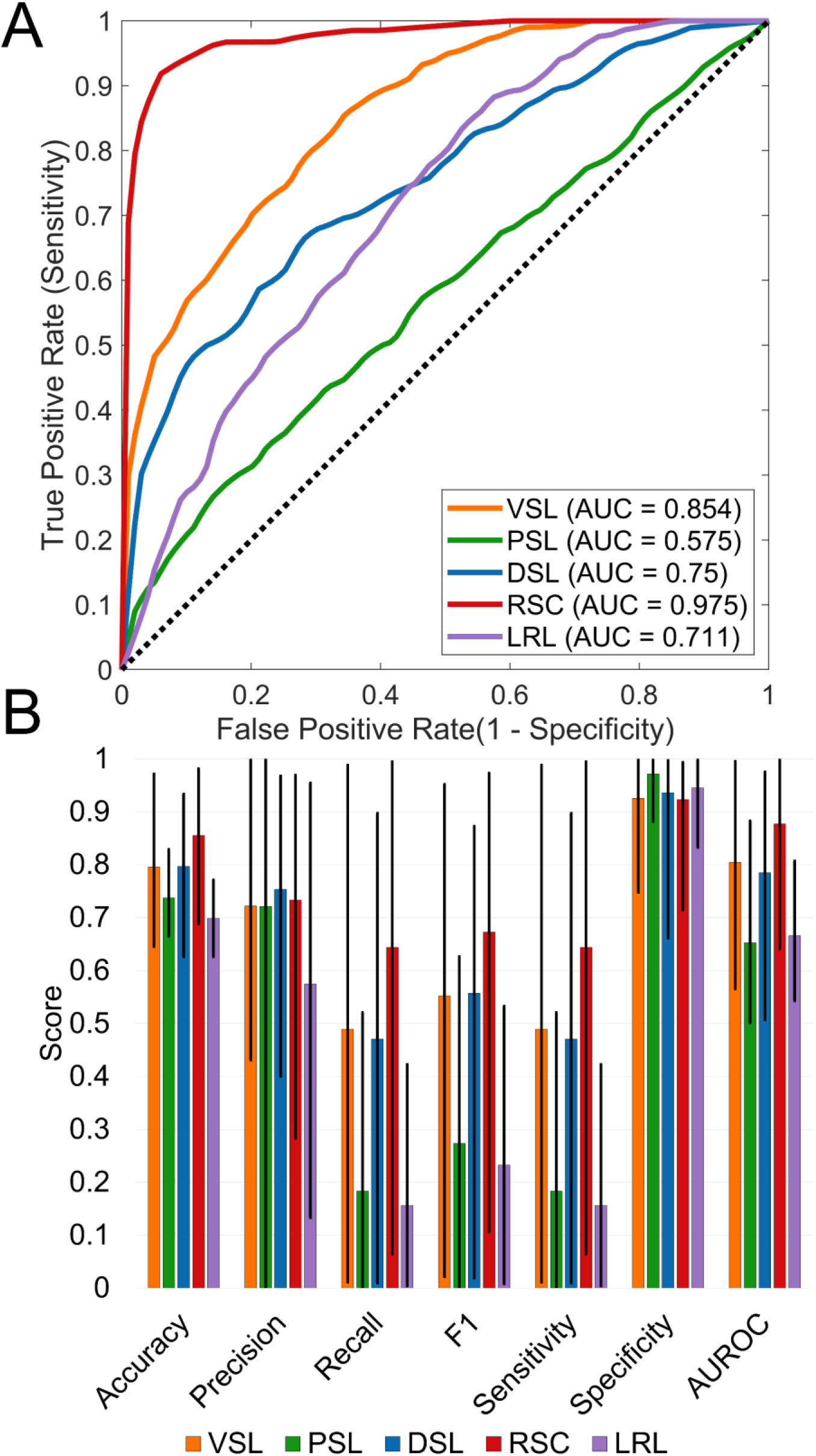
Trained AI model classification performance. (A) Representative receiver operating curves (ROC) and area under the curve (AUC) for the AI model from a subset of the data with wrist radial deviation angles between 10° - 15° between the proximal surface of the scaphoid and the radial surface of the capitate. The dashed line represents a pure random classifier, with an AUC of 0.5. (B) AI model performance metrics across all possible bone surface pairs and wrist angle subsets. Bars show the mean and error bars represent the minimum to maximum. For all metrics, a value of 1.0 is considered optimum.

**Table 1:** Diagnostic Performance of the AI Model for Detection of Wrist Ligament Injuries across all possible bone surface pairs and wrist angle subsets. Values are reported as average [minimum, maximum].

| Injury Type | Accuracy | Sensitivity | Specificity | AUROC |
| --- | --- | --- | --- | --- |
| **VSL** | 0.796 [0.645, 0.972] | 0.490 [0.011, 0.988] | 0.926 [0.748, 1.000] | 0.804 [0.566, 0.996] |
| **PSL** | 0.737 [0.665, 0.829] | 0.183 [0.000, 0.521] | 0.972 [0.882, 1.000] | 0.652 [0.501, 0.883] |
| **DSL** | 0.797 [0.626, 0.933] | 0.471 [0.010, 0.897] | 0.936 [0.661, 0.997] | 0.785 [0.507, 0.975] |
| **RSC** | 0.855 [0.688, 0.982] | 0.644 [0.064, 0.995] | 0.923 [0.715, 0.993] | 0.877 [0.641, 0.999] |
| **LRL** | 0.699 [0.626, 0.771] | 0.156 [0.004, 0.423] | 0.945 [0.833, 0.999] | 0.666 [0.543, 0.807] |

When using a subset of data selected over a clinically- and functionally relevant wrist angle range, namely all images with wrist radial deviation angles between 10 - 15°, the accuracy increased to 0.804 with average AUROC of 0.803 for the five ligaments. Similarly, using a subset of data for all wrist extension angles between 20 - 30°, the accuracy was 0.814, and the average AUROC was 0.824. Generally, the highest classification accuracies were found in images with the wrist in extension or radial deviation.

Overall, the RSC was most accurately classified as intact or injured; individual combinations of bone surface pairs and wrist angle subsets yielded average and maximum AUROCs of 0.877 and 0.999. The LRL was the least accurately classified, with average and maximum AUROCs of 0.666 and 0.807 (Figure 5B). In every ligament, the specificity was greater than the sensitivity. Average specificity exceeded 0.900 for each ligament; mean sensitivity was less than 0.650. Still, certain subsets of bone surface pairs and wrist angles yielded sensitivities above 0.875, with the DSL, VSL, and RSC achieving maximum sensitivities of 0.897, 0.988 and 0.995, respectively (Figure 5B). For example, for the VSL this was the image of the proximal surfaces of the scaphoid and lunate at a slight radial deviation (10 - 15°).

## Discussion

Despite advances in cross-sectional and dynamic imaging (e.g., fluoroscopy, ultrasound, dynamic CT), clinical assessment of wrist ligament injuries remains suboptimal, particularly in detecting subtle or functionally significant instability (6, 7). Even with sophisticated imaging modalities, structural findings do not always correlate with the underlying biomechanical conditions of ligament disruption. Although arthroscopy is regarded as the diagnostic reference standard, surgeons need objective, biomechanics-informed, and comprehensive assessment tools to reduce reliance on invasive procedures. The primary clinical motivation of this study is to provide new tools for clinicians to understand the likelihood of injury to specific scaphoid-lunate ligament stabilizers.

Compared to medical imaging alone, FEMs can provide additional insights into the underlying joint mechanics, that may be beneficial for improving diagnosis of injuries and may even enable targeted treatment strategies. Still, FEM remains slow. In contrast, AI algorithms can make predictions much faster than FEM, and given enough training examples, can often make highly accurate predictions. However, data of musculoskeletal injuries, particularly for subtle injuries, to train these models is scarce. This study demonstrates the feasibility of a high-throughput computational pipeline harnessing synthetic images from FEMs to empower AI models in precisely classifying complex wrist ligament injuries.

Using 4DCT data from two participants, we generated 7,500 biomechanical simulations to bridge musculoskeletal biomechanics with AI. The results suggest that FEM-derived synthetic data can help overcome imaging data scarcity, which limits the application of AI in musculoskeletal research.

Our approach offers a key advantage over traditional AI training methods based on *in vivo* or *in vitro* data alone, namely the FEM-derived synthetic datasets contain simulated ligament injury patterns that are uncommon in clinical or laboratory settings. While these patterns may be clinically unimportant due to their limited prevalence, they can nonetheless provide valuable information to AI models regarding the individual, often subtle, roles of different SL stabilizers, informed by foundational mechanical principles. This enables models to better identify the unique, sometimes subtle, changes resulting from injury to each ligament, thereby enhancing overall performance.

A particular novelty of this work was in how the training data were provided for the AI models. Complex spatiotemporal articular interactions, in the form of proximity maps, were converted to RGB images, which have significantly greater prevalence in AI. In our algorithm, we provided additional descriptive metrics as supplementary inputs to provide contextual information, which improved performance in our initial testing. This provides a promising pathway to combine medical imaging or FEM-derived synthetic data with contextual, and oftentimes crucial, clinical information.

Simulated ligamentous injuries to the DSL, VSL, and RSC were more discernable by the AI model, with maximum specificities and sensitivities all exceeding 0.875 for certain bone surface pairs and wrist angles. In contrast, injuries to the PSL and LRL consistently yielded poorer classification performance. The altered kinematics provided by the DSL, VSL, and RSC are potentially more mechanically consequential for joint surface interactions compared with PSL or LRL. While the model’s overall average AUROC of 0.757 is promising and comparable to prior work (20), performance increased notably when analyzing subsets of data in 10°-15° radial deviation (AUROC of 0.803) and wrist extension (AUROC of 0.824). This aligns with prior work that also showed that wrist extension yielded the greatest discrimination (AUROC of 0.795) (20), highlighting the importance of evaluating clinically-relevant wrist angle ranges. Extension aligns with the common injury mechanism of falling onto an outstretched hand (21), while radial deviation mirrors the clinical stress manipulation used to reveal underlying instability (36). Furthermore, focusing the AI on the relevant bone interactions, the specific articulating surfaces directly governed by the injured ligament, maximizes the discriminative signal. Therefore, enhanced accuracy in these specific subsets likely reflects a biomechanically meaningful amplification of instability rather than selective optimization.

Overall, while the AI models achieved high performance in classifying certain ligaments, at specific wrist angle ranges, and specific image subsets, the results demonstrated substantial variability. While model specificity remained high across all ligaments, sensitivity was more variable and exceeded 0.90 only for selected ligaments and image subsets. While high overall specificity is important for stratifying cases wherein surgical interventions are appropriate, the relatively low sensitivities for non-extension and non-radial deviation positions highlight areas of potential improvement. Future work will focus on identifying the optimal functional ranges for assessing alignment changes at each joint surface from injury. This approach aims to maximize diagnostic capability, providing clinicians with a more nuanced and accurate method for diagnosing specific ligament injuries.

While the overall performance achievements were moderate, this work demonstrates a novel method to synthetically generate data of the hand from computational models to be used for training AI models to diagnose wrist ligament injuries. This is likely amongst the first studies to have ever used these techniques to improve the diagnosis of soft-tissue injuries of the wrist. However, these achievements should be viewed considering certain limitations that affected this work.

First, the synthetically-generated data were derived from two asymptomatic participants, and no experimental data was used to train the AI models. This work attempted to overcome the data scarcity that conventionally limits traditional AI work by generating 9,000,000 physics-driven synthetic images for data augmentation. While our initial results demonstrate feasibility of such methods, future work should involve larger cohorts to maximize generalizability and test models against experimental data of individuals, including patients with arthroscopically confirmed SLIL injuries, not used to create the models.

Second, validation remains a critical concern in AI. Models trained on FEM-derived synthetic data inevitably propagate inherent uncertainties and simplifications from the original FEMs. The scarcity of real-world data further complicates the issue. Real-world data is high-dimensional and has inherent errors that are difficult to replicate in digital data. While simplifications are necessary to efficiently develop models, the impacts on AI model performance remain unknown. While ongoing work in knee modeling has aimed to investigate such questions (37, 38), similar applications to the wrist is limited.

Third, one limitation is the class imbalance introduced by assigning a fixed ⅓ independent injury probability to each ligament. Although this approach effectively models real-world clinical prevalence, it generates uneven training permutations, contrasting rare five-ligament injuries with more common isolated tears. Because deep learning models generally achieve higher robustness with balanced training data, this natural imbalance may have limited overall performance. Future work will leverage the flexibility of synthetic data to investigate how artificially balancing these training distributions impacts diagnostic accuracy.

Lastly, while FEM simulations rely on fundamental mechanical principles that can be physically interpreted, converting these simulation results into RGB vector fields, and then into AI models, significantly reduces the interpretability of the model predictions. While methods like GradCam may help improve the explainability of models (39), they are limited in how much can be gained. Recent developments in the AI field, such as physics-informed neural networks (PINNs) (40), have aimed to improve model interpretability by forcing models to learn underlying mechanics. Future work should integrate physics-informed methods to improve model interpretability and ensure that their predictions hold with well-known biomechanical patterns.

## Conclusion

This study demonstrates that AI models trained on synthetically generated FEM data can classify wrist ligament injuries with similar or better performance than reported previously with other methods and that may be clinically meaningful. We established a scalable framework that bridges computational modeling and data-driven classification to detect various wrist ligament injuries. We developed new approaches to convert complex joint proximity maps into images that are more compatible with AI algorithms. Additionally, we demonstrated techniques for integrating imaging data with clinical metrics to enhance the performance of AI models in future applications. Lastly, the high variability in model performance highlights the importance of utilizing clinically-relevant functional joint ranges, and bone interactions. Future work will focus on validation using patient-specific and clinically acquired datasets, extending to full range of motion and other joint interfaces, improving feature representations to enhance sensitivity, and integrating into clinical platforms to assess its diagnostic impact in the real world.

## Acknowledgements

This manuscript is the result of funding in whole or in part by the National Institutes of Health (NIH) through grants T32 AR056950, F31 AR082227, R01 AR071338, T32 GM065841, and T32 GM145408. It is subject to the NIH Public Access Policy. Through acceptance of this federal funding, NIH has been given a right to make this manuscript publicly available in PubMed Central upon the Official Date of Publication, as defined by NIH. Furthermore, this work was supported by the Biomedical Imaging Resource (BIR) Core Facility at Mayo Clinic, with funding from the Mayo Clinic Office of Core Shared Services through the Early-Stage Investigator Research Award.

## Notes

https://doi.org/10.5281/zenodo.18166610

### Competing Interest Statement

The authors have declared no competing interest.

## References

1. Chan JJ, Xiao RC, Hasija R, Huang HH, Kim JM. Epidemiology of Hand and Wrist Injuries in Collegiate-Level Athletes in the United States. J Hand Surg Am. 2023;48(3):307.e1-.e7.

2. Fowler TP. Intercarpal Ligament Injuries Associated With Distal Radius Fractures. J Am Acad Orthop Surg. 2019;27(20):e893–e901.

3. O’Brien L, Robinson L, Lim E, O’Sullivan H, Kavnoudias H. Cumulative incidence of carpal instability 12-24 months after fall onto outstretched hand. J Hand Ther. 2018;31(3):282–6.

4. Schmitt R, Reidler P, Haas-Lützenberger E, Ruettermann M, Hesse N. Identification of degenerative precursors at the wrist with advanced imaging: current updates. J Hand Surg Eur Vol. 2025;50(1):15–26.

5. Wessel LE, Wolfe SW. Scapholunate Instability: Diagnosis and Management - Anatomy, Kinematics, and Clinical Assessment - Part I. J Hand Surg Am. 2023;48(11):1139–49.

6. Schmitt R, Froehner S, Coblenz G, Christopoulos G. Carpal instability. Eur Radiol. 2006;16(10):2161–78.

7. Bergh TH, Lindau T, Bernardshaw SV, Behzadi M, Soldal LA, Steen K, et al. A new definition of wrist sprain necessary after findings in a prospective MRI study. Injury. 2012;43(10):1732–42.

8. Schmauss D, Pöhlmann S, Lohmeyer JA, Germann G, Bickert B, Megerle K. Clinical tests and magnetic resonance imaging have limited diagnostic value for triangular fibrocartilaginous complex lesions. Arch Orthop Trauma Surg. 2016;136(6):873–80.

9. Torabi M, Lenchik L, Beaman FD, Wessell DE, Bussell JK, Cassidy RC, et al. ACR Appropriateness Criteria(®) Acute Hand and Wrist Trauma. J Am Coll Radiol. 2019;16(5s):S7–s17.

10. Stensby JD, Fox MG, Nacey N, Blankenbaker DG, Frick MA, Jawetz ST, et al. ACR Appropriateness Criteria® Chronic Hand and Wrist Pain: 2023 Update. J Am Coll Radiol. 2024;21(6s):S65–s78.

11. Andersson JK, Andernord D, Karlsson J, Fridén J. Efficacy of Magnetic Resonance Imaging and Clinical Tests in Diagnostics of Wrist Ligament Injuries: A Systematic Review. Arthroscopy. 2015;31(10):2014–20.e2.

12. Götestrand S, Flondell M, Lundin B, Aksyuk E, Shalhoub RA, Szaro P, et al. MRI of wrist ligament trauma was similar at 7 T and 3 T with arthroscopy as a reference standard. Eur Radiol. 2025;35(11):6949–57.

13. Stirling PHC, Duckworth AD, Adams JE, Kakar S. What is the role of arthroscopy in hand and wrist trauma? Bone Joint J. 2025;107-b(3):291–5.

14. Kastenberger T, Kaiser P, Schmidle G, Schwendinger P, Gabl M, Arora R. Arthroscopic assisted treatment of distal radius fractures and concomitant injuries. Arch Orthop Trauma Surg. 2020;140(5):623–38.

15. Henry MH. Management of acute triangular fibrocartilage complex injury of the wrist. J Am Acad Orthop Surg. 2008;16(6):320–9.

16. Nauwelaers M, El Amiri L, Bierry G, Willaume T. Diagnostic accuracy of wrist MR-arthrography protocol using iodinated contrast agent only in the detection of intrinsic wrist ligament and triangular fibrocartilage complex tears. Skeletal Radiol. 2026;55(1):29–38.

17. Erdemir A, Besier TF, Halloran JP, Imhauser CW, Laz PJ, Morrison TM, et al. Deciphering the "Art" in Modeling and Simulation of the Knee Joint: Overall Strategy. J Biomech Eng. 2019;141(7):0710021–07100210.

18. Henak CR, Anderson AE, Weiss JA. Subject-specific analysis of joint contact mechanics: application to the study of osteoarthritis and surgical planning. J Biomech Eng. 2013;135(2):021003.

19. Modaresi S, Kallem MS, Lee P, McIff TE, Toby EB, Fischer KJ. Evaluation of midcarpal capitate contact mechanics in normal, injured and post-operative wrists. Clin Biomech (Bristol). 2017;47:96–102.

20. Trentadue TP, Thoreson AR, Lopez C, Breighner RE, An KN, Holmes DR, 3rd, et al. Detection of scapholunate interosseous ligament injury using dynamic computed tomography-derived arthrokinematics: A prospective clinical trial. Med Eng Phys. 2024;128:104172.

21. Trentadue TP, Lopez C, Breighner RE, Akbari-Shandiz M, An KN, Leng S, et al. Assessing carpal kinematics following scapholunate interosseous ligament injury ex vivo using four-dimensional dynamic computed tomography. Clin Biomech (Bristol). 2023;107:106007.

22. Andreassen TE, Trentadue TP, Thoreson AR, An K-N, Kakar S, Zhao KD. Rapid Development of Efficient Participant-Specific Computational Models of the Wrist. arXiv preprint arXiv:250519282. 2025.

23. Andreassen TE, Trentadue TP, Thoreson AR, Vilai P, An K-N, Kakar S, et al. Finite Element Modeling of the Scaphoid Shift Maneuver: Implications for Scapholunate Ligament injuries. bioRxiv. 2026:2026.02.17.705556.

24. Cooper RJ, Wilcox RK, Jones AC. Finite element models of the tibiofemoral joint: A review of validation approaches and modelling challenges. Med Eng Phys. 2019;74:1–12.

25. Lesage R, Van Oudheusden M, Schievano S, Van Hoyweghen I, Geris L, Capelli C. Mapping the use of computational modelling and simulation in clinics: A survey. Frontiers in Medical Technology. 2023;5:1125524.

26. Tappan I, Lindbeck EM, Nichols JA, Harley JB. Explainable AI Elucidates Musculoskeletal Biomechanics: A Case Study Using Wrist Surgeries. Ann Biomed Eng. 2024;52(3):498–509.

27. Chishti S, Jaggi KR, Saini A, Agarwal G, Ranjan A. Artificial Intelligence-Based Differential Diagnosis: Development and Validation of a Probabilistic Model to Address Lack of Large-Scale Clinical Datasets. J Med Internet Res. 2020;22(4):e17550.

28. Trentadue TP, Thoreson A, Lopez C, Breighner RE, Leng S, Holmes III DR, et al. Morphology of the scaphotrapeziotrapezoid joint: A multi-domain statistical shape modeling approach. Journal of Orthopaedic Research®. 2024;42(11):2562–74.

29. Trentadue TP, Thoreson A, Lopez C, Breighner RE, Leng S, Kakar S, et al. Sex differences in photon-counting detector computed tomography-derived scaphotrapeziotrapezoid joint morphometrics. Skeletal Radiology. 2025:1–11.

30. Zhao K, Breighner R, Holmes III D, Leng S, McCollough C, An K-N. A technique for quantifying wrist motion using four-dimensional computed tomography: approach and validation. Journal of biomechanical engineering. 2015;137(7):074501.

31. Andreassen TE, Hume DR, Hamilton LD, Higinbotham SE, Shelburne KB. Automated 2D and 3D finite element overclosure adjustment and mesh morphing using generalized regression neural networks. Medical engineering & physics. 2024;126:104136.

32. Neu CP, Crisco JJ, Wolfe SW. In vivo kinematic behavior of the radio-capitate joint during wrist flexion-extension and radio-ulnar deviation. J Biomech. 2001;34(11):1429–38.

33. Choi PT, Lui LM. Fast disk conformal parameterization of simply-connected open surfaces. Journal of Scientific Computing. 2015;65(3):1065–90.

34. Abadi M, Barham P, Chen J, Chen Z, Davis A, Dean J, et al., editors. {TensorFlow}: a system for {Large-Scale} machine learning. 12th USENIX symposium on operating systems design and implementation (OSDI 16); 2016.

35. Kingma DP, Ba J. Adam: A method for stochastic optimization. arXiv preprint arXiv:14126980. 2014.

36. Ozçelik A, Günal I, Köse N. Stress views in the radiography of scapholunate instability. Eur J Radiol. 2005;56(3):358–61.

37. Andreassen TE, Hume DR, Hamilton LD, Hegg SL, Higinbotham SE, Shelburne KB. Validating subject-specific knee models from in vivo measurements. Frontiers in bioengineering and biotechnology. 2025;13:1554836.

38. Nazem M, Andreassen T, Kim N, Moyle K, Besier TF, Halloran JP, et al. Deciphering the “Art” in Modeling and Simulation of the Knee Joint: Model Benchmarking. Journal of Biomechanical Engineering. 2026:1–29.

39. Selvaraju RR, Cogswell M, Das A, Vedantam R, Parikh D, Batra D, editors. Grad-cam: Visual explanations from deep networks via gradient-based localization. Proceedings of the IEEE international conference on computer vision; 2017.

40. Raissi M, Perdikaris P, Karniadakis GE. Physics-informed neural networks: A deep learning framework for solving forward and inverse problems involving nonlinear partial differential equations. Journal of Computational physics. 2019;378:686–707.

